# Influence of chemically and biosynthesized silver nanoparticles on in vitro viability and infectivity of Trichinella spiralis muscle larvae

**DOI:** 10.1101/2020.12.17.423206

**Authors:** Salwa Mahmoud Abd-ELrahman, Ahmed Kamal Dyab, Abeer El-sayed Mahmoud, Nahed Ahmed Elossily, Fahd M. Alsharif, Shaymaa M. Mohamed, Mosleh Mohammed Abomughaid

**Affiliations:** Department of Parasitology, Faculty of Veterinary Medicine, Assiut University, Egypt; Department of Medical Parasitology Faculty of Medicine, Assiut University, Egypt; Department of Pharmacognosy, Faculty of Pharmacy, Assiut University, Egypt; Department of industrial pharmacy department, Faculty of pharmacy, Al Azhar University, Assiut; Department of Medical laboratory Sciences University of Bisha, Bisha, Saudia Arabia

## Abstract

**Back ground:** Trichinellosis is a serious worldwide parasitic zoonosis. The available therapy for the treatment of ***Trichinella spiralis*** is not satisfactory. This work aimed at evaluating of the in vitro effect of silver Therefore, the recovery of effective treatment is required.nanoparticles (AgNPs) on muscle larvae of ***Trichinella***.

**Methodology / principal finding:** The present study investigated the larvicidal properties of chemical and myrrh AgNPs on muscle larvae (ML) of ***T. spiralis***. The used AgNPs were chemically prepared using NaBH4 as reducing agent and biosynthesized using methanolic myrrh extract. Characterization of synthesized AgNPs was monitored via UV-Vis spectrophotometry, Fourier transform infrared spectroscopy and transmission electron microscopy (TEM) studies. The ML incubated with AgNPs at concentrations ranged from 1μg/ml to 20μg/ml.

**Conclusions /Significance:** Chemical and biosynthesized AgNPs revealed marked larvicidal effect against ML of ***Trichinella***. Additionally, this *in vitro* study showed degenerative changes affecting the cuticle of AgNPs treated ML. The effectiveness of AgNPs on the infectivity of ***Trichinella*** ML was also assessed. The results showed complete inhibition of the infectivity of ML exposed to sublethal doses of chemical and myrrh prepared AgNPs when used to infect animal models. This is the first report where myrrh synthesized AgNPs have been tested for their anthelminthic activity against ***Trichinella*** in an *in vitro* model.

**Author summary:** Trichinellosis is a serious worldwide parasitic zoonosis. The available therapy for the treatment of ***Trichinella spiralis*** is not satisfactory. Therefore, the recovery of effective treatment is required. The present study investigated the larvicidal properties of chemical and myrrh AgNPs on muscle larvae (ML) of ***T. spiralis***. The ML incubated with AgNPs at concentrations ranged from 1μg/ml to 20μg/ml. Chemical and biosynthesized AgNPs revealed marked larvicidal effect against ML of ***Trichinella***. Additionally, this *in vitro* study showed degenerative changes affecting the cuticle of AgNPs treated ML. Also the results showed complete inhibition of the infectivity of ML exposed to sublethal doses of chemical and myrrh prepared AgNPs when used to infect animal models.

## Introduction

Trichinellosis is a global serious zoonotic parasitic disease induced by the genus ***Trichinella***. Different developmental stages of ***Trichinella spiralis***, adult, migratory and encysted larvae are found in the same host and affects a wide diversity of mammal as well as human. so this parasite has been commonly used as an experimental model to evaluate the effect of many anthelmintic agents (1). Human infections occur mainly through the consumption of infective larvae found in undercooked pork meat products (2). Benzimidazole derivatives such as albendazole, flubendazole and mebendazole are the main anthelmintic drugs used for the treatment of trichinellosis (3). Nevertheless, they have restricted bioavailability with the risk of resistance and weak activity against encapsulated muscle larvae (4). Additionally, some of these drugs are contraindicated in pregnant women and children under three years old (1), while others are supposed to be carcinogenic (5). Consequently, efforts to recover safe and effective antitrichinellosis drugs are required, especially those derived from medicinal plants (1), as they have less toxicity and are free from adverse effects (6).

Myrrh, resinous exudate,derived from the stem of ***Commiphora species*** and comprises gum, volatile oil and resin. It is one of the most commonly used plant exudates in traditional medicine as it retains antipyretic, analgesic, antibacterial and antifungal properties (7). Moreover, Myrrh possesses fasciolocidal, schistosomicidal, insecticidal, molluscicidal activities and an effective activity against both intestinal and muscular stages of ***T. spiralis*** (6).

Synthesis of metal nanoparticles (NPs) is a growing research area due to the extensive applications in several fields such as medicine, electronics, and energy (8). Noble metals such as gold (Au), silver (Ag) and platinum (Pt) are considered as the most important classes of metal NPs (9). Among the noble metal NPs, AgNPs have emerged as one of the fastest growing materials owing to their unique physical, chemical and biological properties. The small sized particles and the large surface area of AgNPs are the main features responsible to their potent antimicrobial activity against a wide range of pathogenic microorganisms (10).

The production of significant amounts of AgNPs is attained with several physical and chemical techniques comprising laser ablation, lithography and the photochemical reduction (11). Nevertheless, synthesis techniques remain relatively expensive and sometimes require the consumption of some hazardous moieties (12). Biosynthesis of AgNPs using biological sources such as bacteria, fungi(13), yeast (14) and plants has been investigated for the green synthesis of AgNPs due to the major advantage of this synthesis procedure in protecting the environment (eco-friendly) (15). Several plant species have been used in this regard, however, only few available reports concerning the antiparasitic activity of metal nanoparticles have been. For example, Marimuthu et al., studied the efficiency of AgNPs against larvae of ***Anopheles subpictus, Rhipicephalus microplus*** and ***Culex quinquefasciatus*** (16). Said et al. validated the antigiardial activity of silver, chitosan and curcumin nanoparticles (17). Lately, Kar, et al. (2014) studied the effectiveness of gold nanoparticles derived from a ***Nigrospora oryzae*** fungus against ***Raillietina*** sp.(19) However, to the best of our knowledge, no reports related to *in vitro* AgNPs anthelminthic effect against ***T. spiralis*** larvae are accessible.

In this study, new series of stable densely discrete AgNPs were produced via a green synthesis approach using the natural non-toxic extract of myrrh as well as the common chemical approach. Myrrh extract has been employed as reducing and capping agent for the synthesis of AgNPs.

. In addition, *in vitro* anthelmintic properties of chemical and biosynthesized AgNPs on ***Trichinella spiralis*** muscle larvae have been investigated.

## Material and methods

### Chemicals

Silver nitrate (AgNO_3_) and sodium borohydride (NaBH_4_, 99.99%) were purchased from Sigma-Aldrich (MO, USA). Polyvinyl pyrrolidone (PVP, M.wt 25K) was obtained from Fisher Scientific (NJ, USA).The myrrh used in this study was purchased from a local commercial store in Assiut city, Egypt.

The strain of *Trichinella spirallis* isolated from a naturally infected pig were obtained from El-Bassatine Abattoir, Cairo, Egypt.

## Methods

### Infectivity of mice with *Trichinella spiralis*

*Trichinella spirallis* strains were maintained by consecutive *in vivo* passages in BALB/c mice at the animal house of Assiut University (Assiut, Egypt) under specific pathogen-free conditions in accordance with the institutional and national guidelines. Animals were fed on a standard diet and tap water. Stool examination of the mice was performed prior to the study to ensure the absence of any possible parasitic infection. Mice were orally infected with 300 ***T. spiralis*** larvae/mouse (20). Larvae were obtained by enzymatic digestion of skeletal muscle from mice 30 to 90 days post infection (pi) as previously described (21).

### Preparation of myrrh methanolic extract

The myrrh used in this study was purchased from a local commercial store in Assiut city, Egypt. It is the oleo-gum resin exudate of (*Commiphora myrrha*) imported from Somalia. Its identity was confirmed through examining its organoleptic and chemical properties(22). A specific weight (250 g) was pulverized to fine powder and macerated in 1 L of methanol for 24 h. Ultrasonication for 1 h was applied, to speed up the extraction process and enhance the yield thereof, then the attained extract was filtered. The aforementioned procedure was repeated five times. All methanolic extracts were collected together and evaporated by means of a rotary evaporator under reduced pressure at 40 °C to obtain a dried residue (59.5 g) representing the alcohol-soluble portion of the used exudate (namely, the essential oil and the alcohol-soluble resins).

### Green synthesis (Biosynthesis) of silver nanoparticles (AgNPs) using *Myrrh* extracts

Silver nanoparticles (AgNPs) at a concentration of 100 µg/mL were prepared via green biosynthesis approach as previously described with some modification (23). Briefly, in a round bottom flask, 0.95 mM AgNO_3_ solution in 20 mL distilled water was added to 20 mL of the previously prepared *Myrrh* extract. The mixture was vigorously stirred on a magnetic stirrer with a hot plate for one hour. The formation of AgNPs was indicated by changing of interaction solution into yellow and then green.

### Chemical synthesis of AgNPs using sodium borohydride (NaBH4)

AgNPs were synthesized, at a concentration of 100 µg/ml, according to previously reported method (24). In brief, 4 mM of freshly prepared sodium borohydride (NaBH4) were dissolved in 20 mL distilled water to which 20 ml aqueous 0.95 mM silver nitrate (AgNO_3_) solution was added in drop wise manner. The mixture was placed in an ice bath and stirred for 30 min. Yellow discoloration of the solution reaction indicating occurrence of reduction interaction with spherical AgNPs formation. Polyvinyl pyrrolidone (PVP) was used as a stabilizer in order to prevent AgNPs aggregation.

### Characterization of AgNPs

The synthesis of AgNPs was confirmed with the help of multiple techniques. The reduction of pure Ag+ ions was confirmed by recording the absorbance (A) on a UV-vis spectrophotometer (Double beam spectrophotometer, UV-160 Shimadzu co., Japan) in a wavelength ranged from 200 to 800 nm (25). In addition, infrared spectra of chemically and biosynthesized AgNPs were recorded using Nicolet iS10 FT-IR Spectrometer (Thermo Fisher Scientific, NJ, USA) over the frequency range 4000-700 cm^-1^ at 4 cm^-1^ resolution. Each sample was prepared by mixing with potassium bromide (KBr) and compressed under hydraulic pressure (25). Further, the size and morphology of both chemically and biosynthesized AgNPs were observed using TEM (JEOL model, JEM-100 CXII, Tokyo, Japan) with an accelerating voltage of 200 kV.

### Antiparasitic activity of AgNPs

Preparation of AgNPs suspension: the suspension of nanoparticles was prepared in five different concentrations (1, 5, 10, 15, 20 μg/ml). Collection of ***T. spiralis*** larvae: muscular larvae were obtained from experimentally infected mice. The larvae were recovered from carcasses of infected mice 30 days pi. by artificial digestion method according to slandered procedures (21) The larvae were washed several times in PBS and diluted in Rapid Prototyping and Manufacturing Institute (RPMI)-1640 medium that containing 10% fetal calf serum and antibiotics (200 U/ml penicillin and 200 μg/ml streptomycin).

Incubation of the parasite with chemically and biosynthesized AgNPs: a total of 1ml of each concentration of AgNPs mentioned above was mixed with 50 μl from larval suspension containing nearly about 100 larvae (counted by light microscope) in 6-well plates which sealed and incubated at 37°C in an atmosphere containing 5% CO^2^ for 1, 4, 24, 48, 72 and 96 h. Untreated larvae incubated with the appropriate volume of PBS and submitted to the same conditions used as controls. There were three replicates for each concentration and control.

At the end of each incubation period, the larvae (both dead and living) in each well were collected, washed with PBS several times to remove adherent particles and counted under stereo-microscope. The viability rates of the parasite were calculated as follows:

Viability rate % = the number of viable parasite/total parasite × 100.

### Electron microscope scanning of the parasite (SEM)

As previously described by Bughdadi, the ultrastructural changes of the treated larvae and controls were investigated by SEM. The larvae were gently washed several times with PBS to remove adherent AgNPs and fixed in 2.5% glutaraldehyde solution for 24h. Then, the fixed larvae re-washed again in PBS for 5 min and post fixed with a solution of 2% osmium tetroxide in sodium cacodylate buffer for 1 h. The specimens were then dehydrated in increasing alcohol concentrations (30%, 50%, 70%, 90% and then 100%) and dried in air then mounted on sputter coated with gold and scanned by SEM (Jeol Jsm-5400, Tokyo, Japan) (26).

### Assessment of the effects on the chemically and biosynthesized AgNPs on the ability of *T. spiralis* larvae to produce infection in experimental animals (27)

A sub-lethal dose of chemically and Myrrh AgNPs were used to detect their effect on the **infectivity** of ***T spiralis*** larvae. Three replicates were prepared by adding larval suspension containing about 300 larvae to 1 ml of 1μg/ml concentration of chemically and biosynthestized nanoparticles. Controls included larvae incubated with the appropriate volume of PBS. The larval suspensions were incubated at 37 C and 5% CO2 for 24 hr. the viability of the larvae and its appearance were observed using a light dissecting microscope. Thereafter, the larvae were collected and used to orally infect 6 Swiss female mice, 8 wk of age for each treated and control groups. The infected mice were sacrificed 5 days pi, their small intestines removed, cut into small pieces and washed to remove intestinal contents. Then, the intestinal pieces were slit longitudinally and the mucosa was gently scraped and incubated in PBS for 4 h at 37 °C. Afterwards, the adult worms were collected and counted by stero-microscope. After 30 days, 3 mice were sacrificed for muscular larvae detection. Regarding the muscular larvae, the other 3 mice were sacrificed 30 days pi for recovery of the larvae.

## Statistical analysis

The collected data were analyzed by Statistical Package for Social Sciences v.20 for Windows (SPSS). The significance of differences between the groups were calculated using the Chi-square test for trend analysis to compare the proportion of viable Larvae in relation to control group (p-value of < 0.001 considered significant). The correlation coefficient (r) was calculated to estimate the degree of similarity in the response of larvae to chemically and biosynthesized AgNPs.

## Results

### Silver nanoparticles synthesis and characterization

A distinct shifting in color to yellow was observed in reaction solution of AgNO_3_ on adding NaBH_4_ indicating the AgNPs formation. At the same time, biosynthesis of AgNPs was detected during the manufacture process via the color change of the myrrh extract from yellow into brown. The obtained UV-vis spectrophotometry spectrum confirmed the formation of the AgNPs as it showed an intense band around 430 nm. This peak is correspondent to surface plasmon resonance (SPR) band which is attributed to the excitation of free electrons in the nanoparticles (Fig. 1). FT-IR spectroscopy analysis was also used to identify the possible functional groups of methanolic *Myrrh* extract involved in the green synthesis of AgNPs. The obtained pattern showed bands around 1445, 1438, 1635, and 3440 cm^-1^. A total disappearance of the peaks at ∼1455 cm^-1^ and ∼1377 cm^-1^ and a slight shift of the peak at ∼1700 cm^-1^ to ∼ 1650 cm^-1^ were observed in the methanolic myrrh extract after the bio-reduction process. In addition to OH broad peak at 3450 cm^-1^, these changes in peaks may prove that polyols and phenols in aqueous myrrh extract were mainly responsible for the reduction of Ag^+^ ions into colloidal Ag (Fig. 2). TEM images of the chemically synthesized AgNPs showed uniform spherical particles in the nano-range (average 20 nm). Likely, TEM micrographs of the biosynthesized AgNPs exhibited particles with uniform shape and in the nano-range (10-25 nm). Besides, the TEM image demonstrated absence of aggregates which suggests the stability of the biosynthesized AgNPs by *myrrh* extract. Moreover, AgNPs morphology was consistent and did not exhibit any changes after 6 months storage at room temperature (Fig 3).

**Fig 1:**
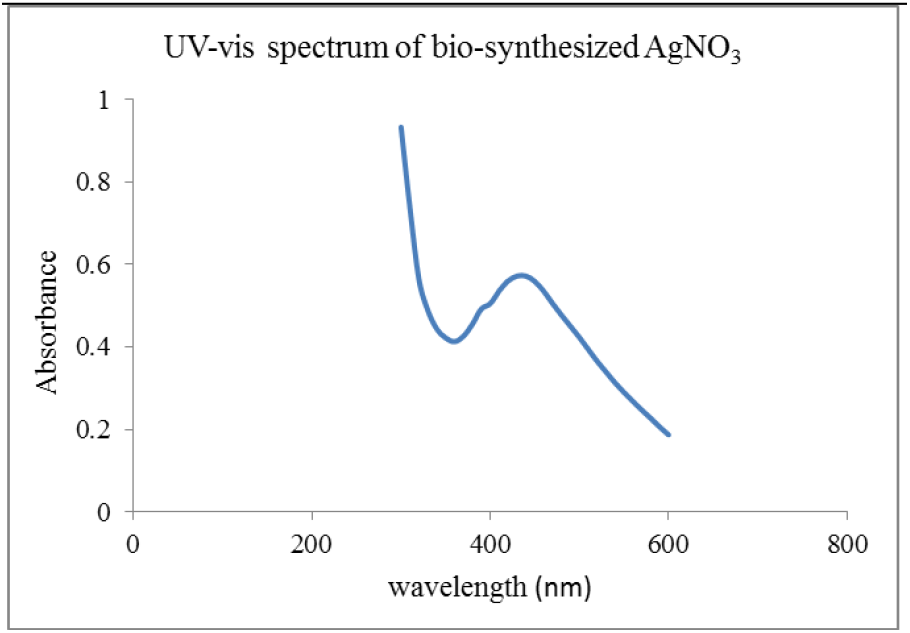
UV-vis spectrum of biosynthesized AgNPs using Myrrh extract

**Fig 2:**
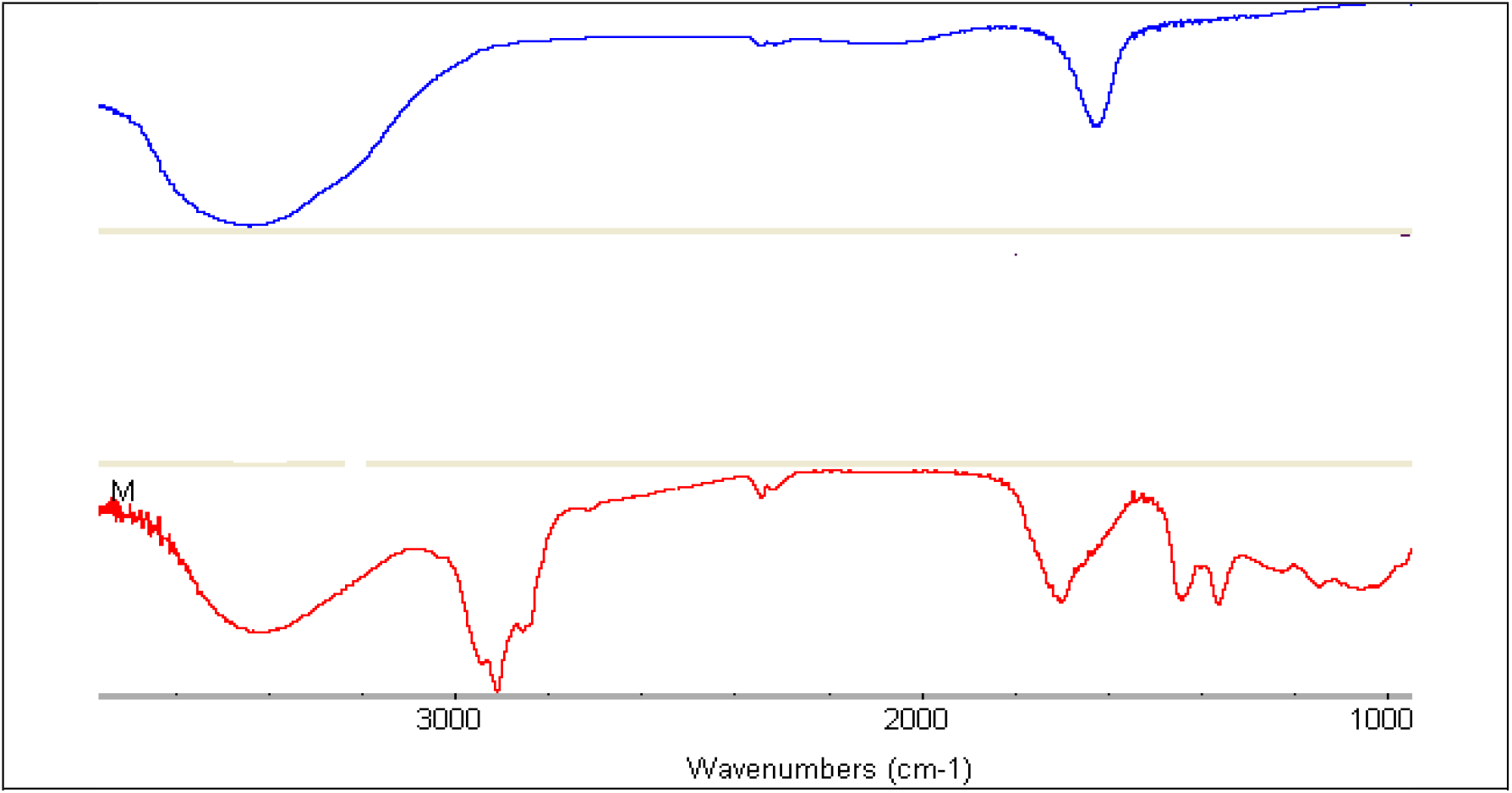
FT-IR spectrum of biosynthesized AgNPs (blue color) *Myrrh* methanolic extract(red color)

**Fig 3:**
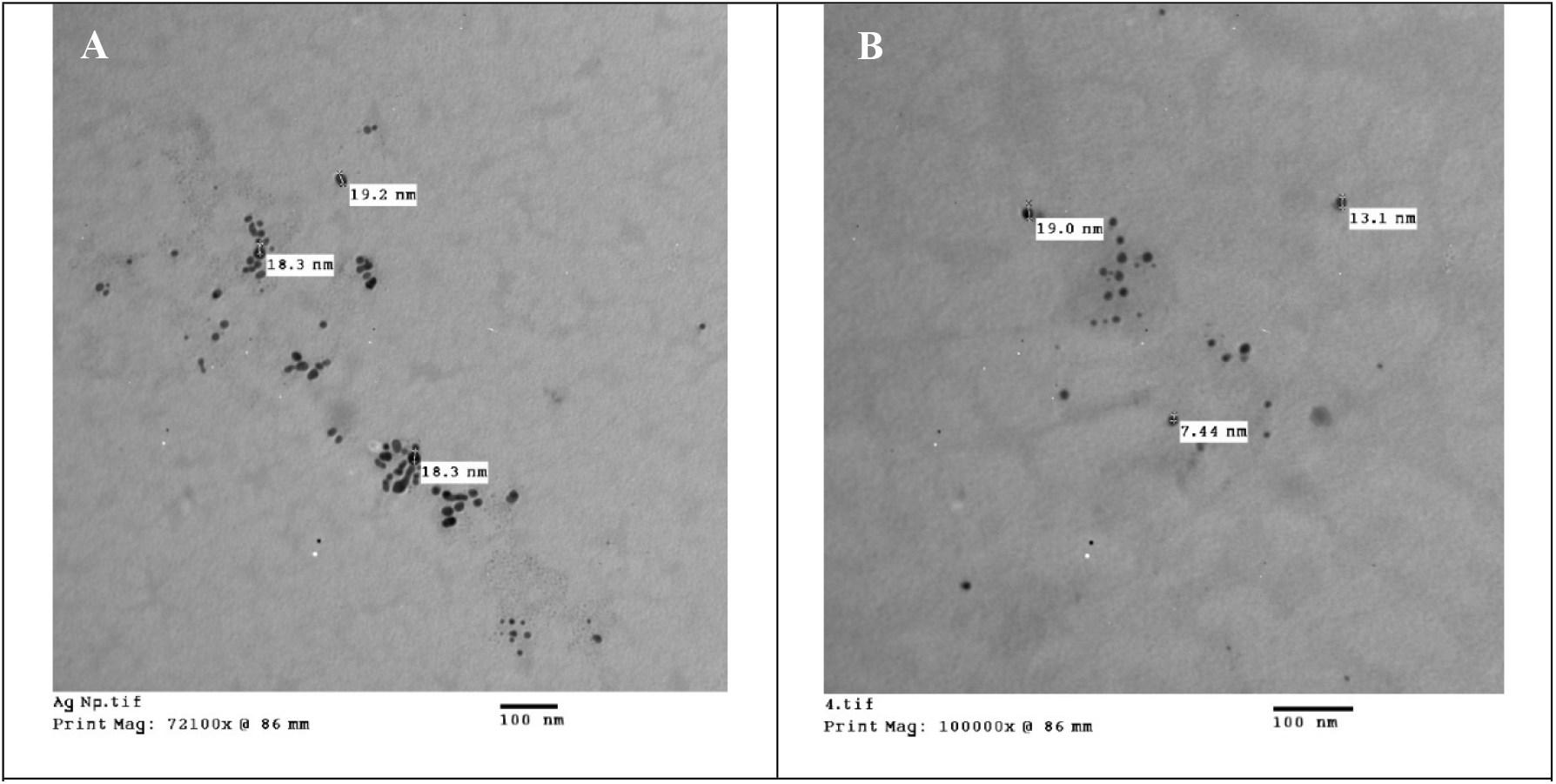
TEM micrographs of chemically (A) and biosynthesized AgNPs (B)

### The *in vitro* effect of chemically and biosynthesized AgNPs on the viability of the muscular larvae of *T. spiralis*

Over the course of AgNPs incubation, the larvae were directly observed under a light microscope; the untreated larvae isolated from striated mouse muscle tissue display the typical coiling behavior of this stage when removed from the nurse cell. On the other hand, the larvae exposed AgNPs showed slow motility, loss of coiling behavior followed by death (Fig 4 A-E).

**Fig 4:**
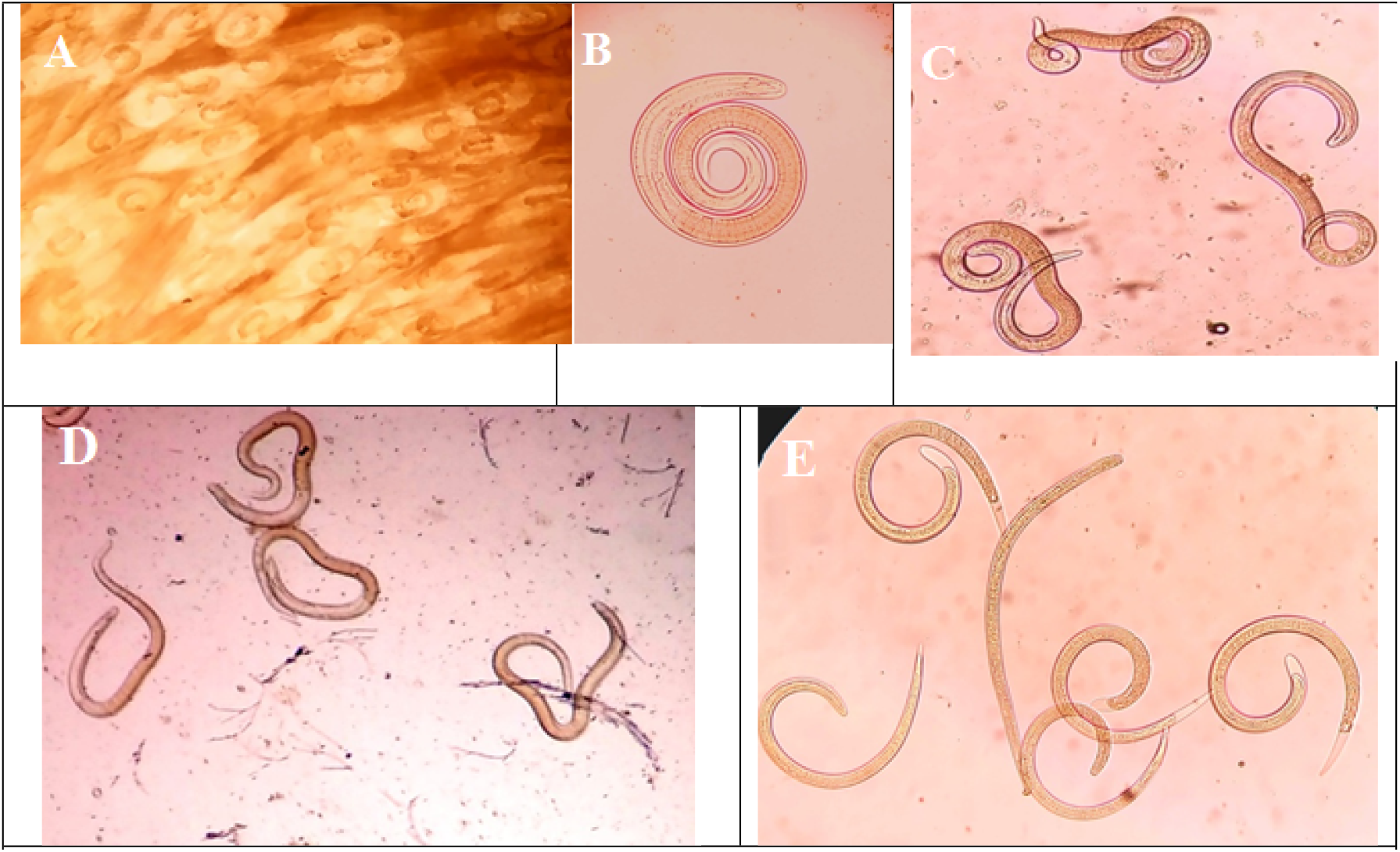
Encysted larvae in muscle of diaphragm x100 (A), Typical coiled larvae x100(B), Active motile larvae x100 (C), Weak larvae after treatment by silver nano particles x 100 (D), Muscular larvae after complete death x 100 (E)

The chemically and biosynthesized nanoparticles showed remarkable effects on the viability of larvae. The larvicidal effect was dose and time dependent, in both AgNPs there was no obvious effect at 1^st^ and 4^th^ h of exposure in all concentrations.

In relation to chemically synthesized AgNPs, a 100% mortality rate was observed after 2 days of exposure at concentrations of 10, 15 and 20 μg/ml, while the viability rates of larva reduced with prolonged exposure at lower concentrations 5 and 1 μg/ml where the death occurred on the 3^rd^ and 4^th^ day of exposure respectively. As compared with controls in, the larvicidal effects of chemically synthesized AgNPs were significantly higher starting from 1st day (P<0.001) (Table 1).

**Table 1:** The Effects of chemically synthesized Silver Nanoparticles on viability of Muscular Larvae of ***Trichinella Spiralis***:

| Dose | 1<sup>st</sup> hour | 4<sup>th</sup> hour | 1<sup>st</sup> day | 2<sup>nd</sup> day | 3<sup>rd</sup> day | 4<sup>th</sup> day | 5<sup>th</sup> day | 6<sup>th</sup> day | 7<sup>th</sup> day | P-value* |
| --- | --- | --- | --- | --- | --- | --- | --- | --- | --- | --- |
| 1µg/ml | 95 | 95 | 71.4 | 47.9 | 20.5 | 0 | 0 | 0 | 0 | < 0.001 |
| 5µg/ml | 95 | 80 | 38.5 | 15.38 | 0 | 0 | 0 | 0 | 0 | < 0.001 |
| 10µg/ml | 95 | 77.5 | 17.14 | 0 | 0 | 0 | 0 | 0 | 0 | < 0.001 |
| 15µg/ml | 95 | 70.33 | 5.3 | 0 | 0 | 0 | 0 | 0 | 0 | < 0.001 |
| 20µg/ml | 95 | 60 | 3.33 | 0 | 0 | 0 | 0 | 0 | 0 | < 0.001 |
| Control group | 95 | 95 | 90 | 85 | 85 | 70 | 50 | 35.29 | 20 | < 0.001 |
| P-value* | =1.000 | =0.925 | < 0.001 | < 0.001 | < 0.001 | < 0.001 | < 0.001 | < 0.001 | < 0.001 |  |

\*Chi-square for trend analysis was used to compare the proportion of viable Larvae.
■ P< 0.001, compared with the corresponding parasite controls. ■ There is decrease in viability rate by time P< 0.001.

Concerning biosynthesized AgNPs, a 100% mortality rate occurred after 2 days of exposure to concentrations of 5, 10, 15 and 20 μg/ml, whereas the viability rates of larva decreased with prolonged exposure at a concentration of 1 μg/ml where the death occurred in third day. There was a significant difference in the mortality rate between all concentrations of green synthesized AgNPs and the control group starting from 1st day (P<0.001) (Table 2).

**Table 2:**
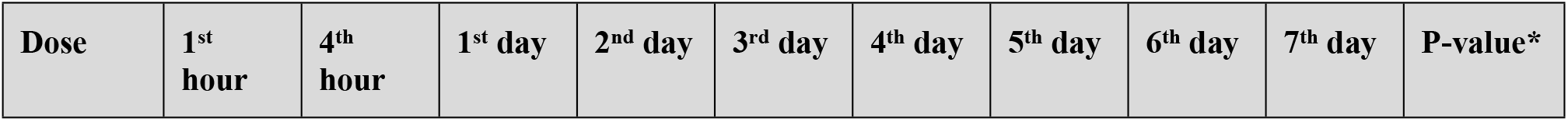
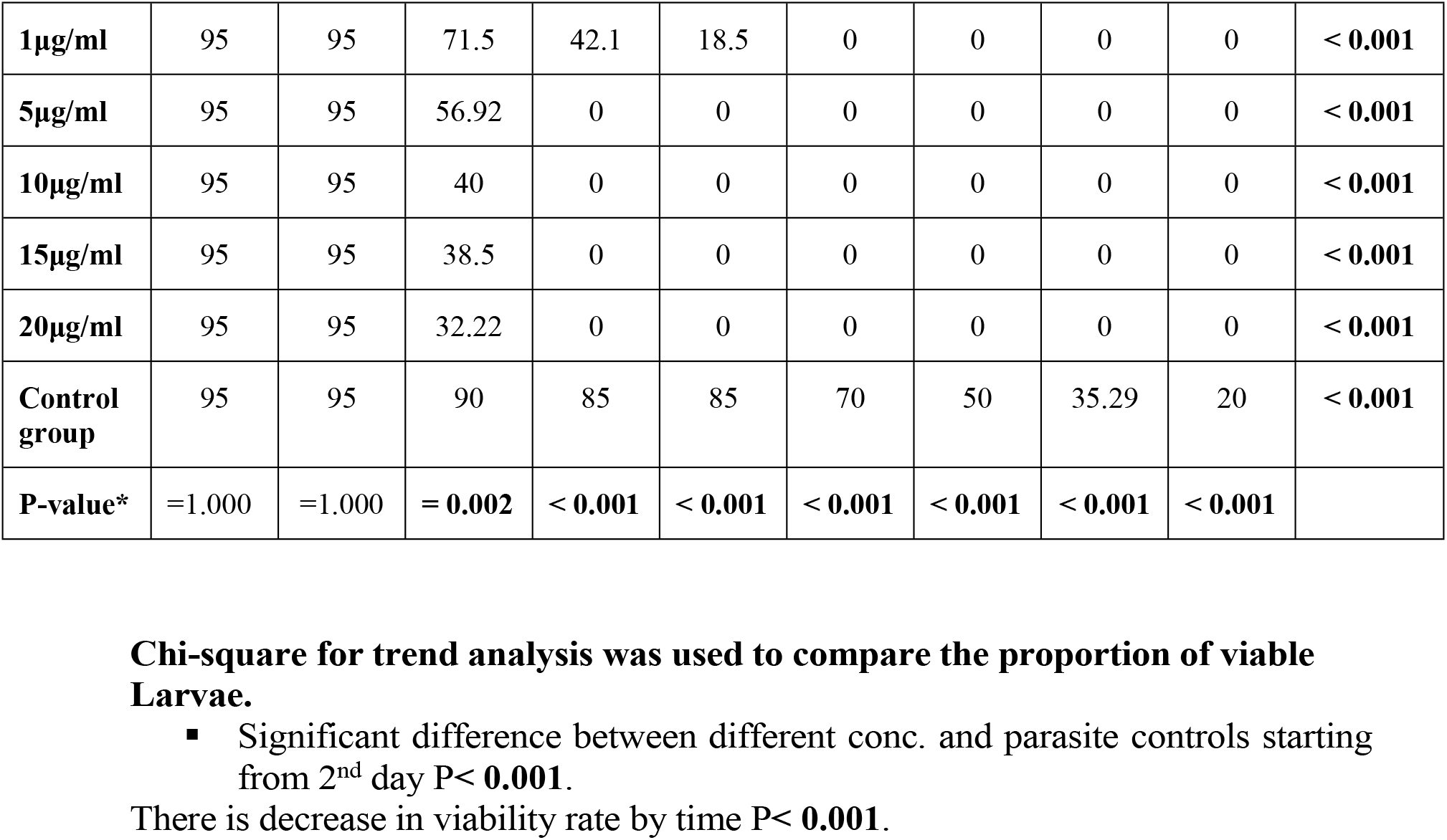
The Effects of Silver Nanoparticles derived from Myrrh on viability of Muscular Larvae of *Trichinella Spiralis*.

| Dose | 1<sup>st</sup> hour | 4<sup>th</sup> hour | 1<sup>st</sup> day | 2<sup>nd</sup> day | 3<sup>rd</sup> day | 4<sup>th</sup> day | 5<sup>th</sup> day | 6<sup>th</sup> day | 7<sup>th</sup> day | P-value* |
| --- | --- | --- | --- | --- | --- | --- | --- | --- | --- | --- |
| 1µg/ml | 95 | 95 | 71.5 | 42.1 | 18.5 | 0 | 0 | 0 | 0 | < 0.001 |
| 5µg/ml | 95 | 95 | 56.92 | 0 | 0 | 0 | 0 | 0 | 0 | < 0.001 |
| 10µg/ml | 95 | 95 | 40 | 0 | 0 | 0 | 0 | 0 | 0 | < 0.001 |
| 15µg/ml | 95 | 95 | 38.5 | 0 | 0 | 0 | 0 | 0 | 0 | < 0.001 |
| 20µg/ml | 95 | 95 | 32.22 | 0 | 0 | 0 | 0 | 0 | 0 | < 0.001 |
| Control group | 95 | 95 | 90 | 85 | 85 | 70 | 50 | 35.29 | 20 | < 0.001 |
| P-value* | =1.000 | =1.000 | = 0.002 | < 0.001 | < 0.001 | < 0.001 | < 0.001 | < 0.001 | < 0.001 |  |

**Chi-square for trend analysis was used to compare the proportion of viable** **Larvae.**
▪ Significant difference between different conc. and parasite controls starting from 2<sup>nd</sup> day P< 0.001.
There is decrease in viability rate by time P< 0.001.

### Effects of AgNPs on the morphology of muscular larvae by SEM

Analysis of the effects of AgNPs on the cuticle of ***T. spiralis*** larvae was conducted using the images obtained by SEM (Fig. 5-7). The results of SEM revealed marked tegmental deformation in larvae exposed to 20 μg/ml of chemically and biosynthesized AgNPs for 24 h when compared to untreated control. The cuticle of the treated larvae turned opaque and showed areas with multiple blebs, vesicles and loss of normal creases. The sloughing of some areas of cuticle was observed. Scanning electron microscope of ***T. spiralis*** larvae of the control group showed normal cuticle with transverse creases and longitudinal ridges.

**Fig 5.**
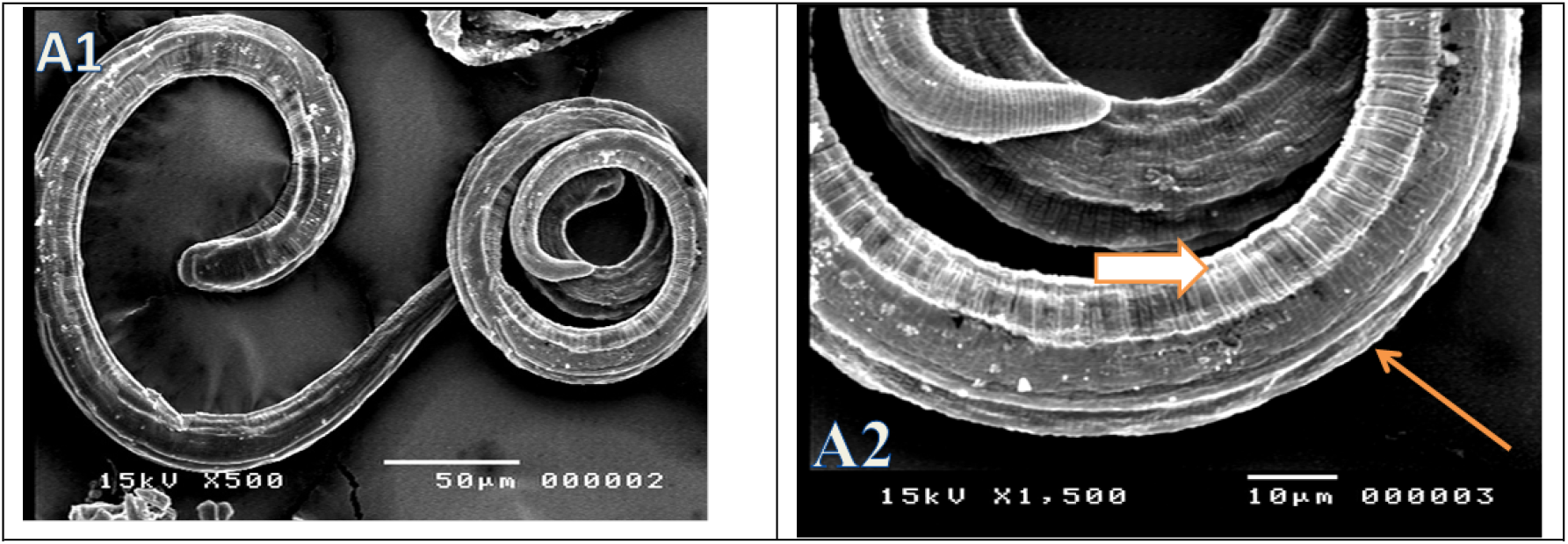
(A1,2): SEM showing normal cuticle of control ML with transverse creases (short arrow) and longitudinal ridges (long arrow)

**Fig 6.**
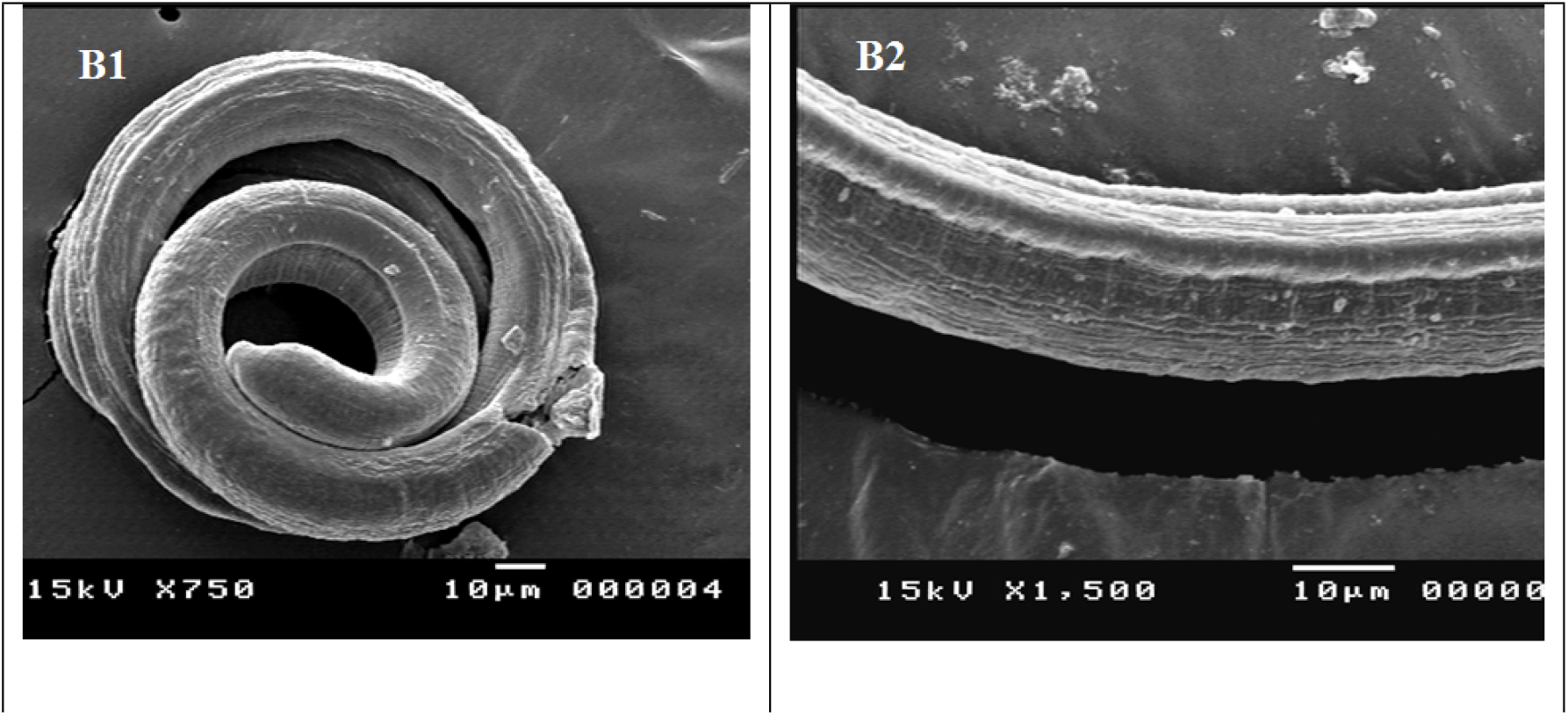
**(B1, 2):** SEM showing ML treated with silver nano particles by SEM showing belbes and vesicles.

**Fig 7:**
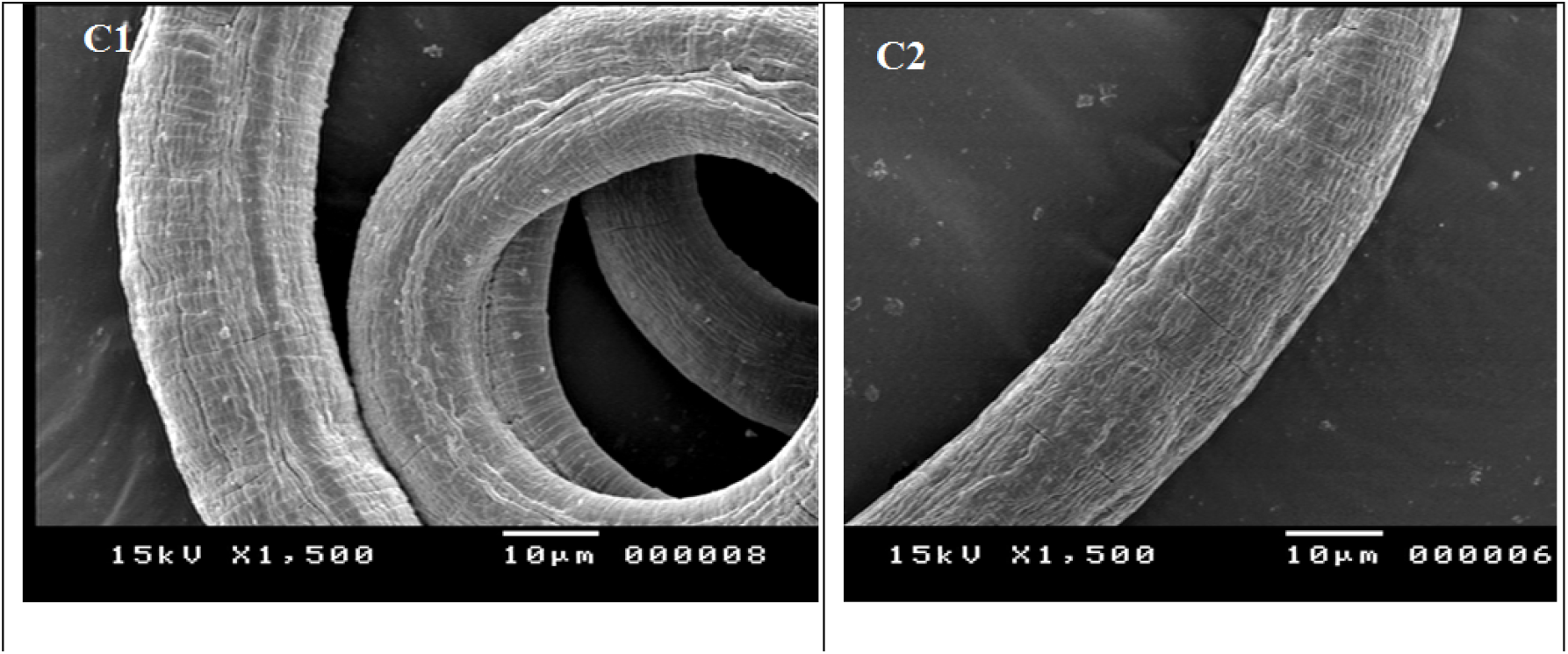
**(C1, 2)**. SEM showing ML treated with silver nano particles prepared with myrrh. Vesicles (long arrow). Belbing (short arrow).

### Effects of AgNPs on the ability of larvae to produce infection in experimental animals

Larvae were exposed to a sub-lethal dose of AgNPs suspension (1μg/ml) for one day. The suspension containing larvae were then used orally to infect mice to study the ability of larvae to develop into an adult (intestinal phase) and to produce muscular larvae (larval phase). Results showed that no adult worms and no muscular larvae were detected in the intestines and muscles of mice infected with treated larvae. While in the control group, examination of the intestine revealed the presence of adults = 228.33 ± 10.3 adult worm. Meanwhile, larvae were also recovered from the muscles mice infected by the control group = 269.3×10^3^ ± 35.9×10^3^. There were significant differences in the infectivity of ***T. spiralis*** larvae treated with AgNPs and control groups (P Value < 0.001).

Based on the obtained data, no significant difference was observed in larvicidal effect between chemically and biosynthesized AgNPs and auspiciously the correlation coefficient between the two treatments was highly positive significant which means that, both treatments have approximately the same lethal effect on ***T. spiralis*** larvae (Fig 8)

**Fig 8:**
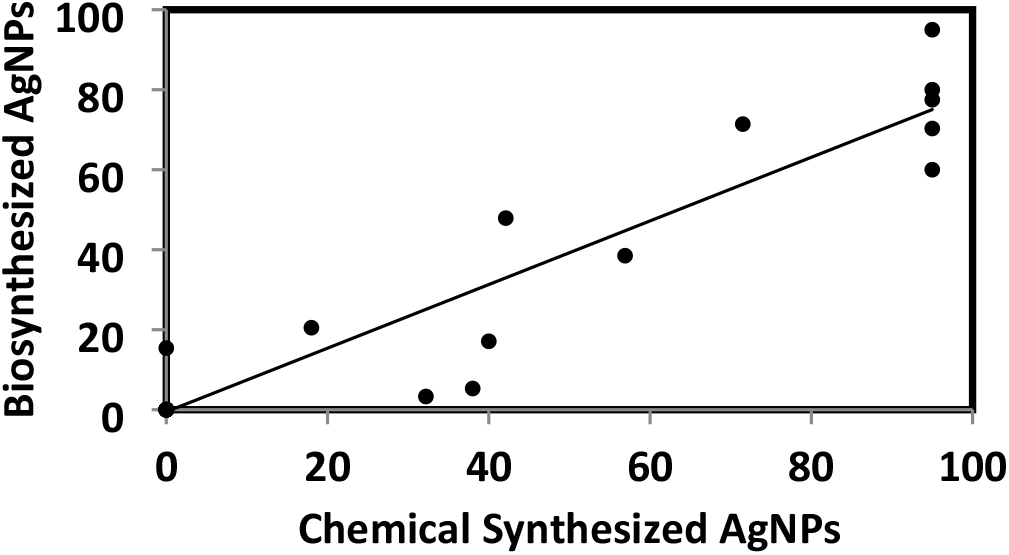
Correlation Coefficient (r) between the effect of chemical and biosynthesized AgNPs on the viability of T. spiralis ML

## Discussion

There is currently a broad consensus that silver nanoparticles have inhibitory effects on many infective agents. Therefore, silver nanoparticles have been used as an alternative treatment against various emerging microbial resistance (28). Myrrh extract has been employed a reducing and capping agent for the synthesis of AgNPs which is more favorable than the microbial synthesis as there is no requirement of the particularized process of culturing and maintaining the cells (23).

The Physicochemical characterization of nanoparticles is essential for better interpretation of results. The nanoparticle characterization of chemically and green synthesized AgNPs was done using a combination of UV-Vis, FT-IR and TEM analysis in order to provide clear insight into crystalline nature, chemical composition, surface property morphology and the particle size. The UV-vis spectroscopy spectrum showed a distinct absorption peak at 430 nm, this might be attributed to the excitation of surface plasmon resonance (SPR) of the synthesized Ag-NPs(29). Many investigators indicated that Ag-NPs exhibited UV-vis absorption spectra which ranged between 410 and 440 nm which were also assigned to other different metal nanoparticles(30), (31).

shift of peak position were observed in FTIR spectrum of both AgNPs also showed intense absorption bands at 1445, 1438, 1635, and 3440 cm^-1^ The band around 1650 cm^-1^ corresponds to aromatic rings. Moreover, the strong broad band appearing at about 3440 cm^-1^ in both FT-IR spectra can be assigned to the stretching vibrations of O-H groups in alcohol and phenols (32). This band assures the presence of several O-H groups in the *myrrh* extract which is anticipated to play a major role in the reduction process of Ag^+^ ions into AgNPs (23). The TEM images of the chemically synthesized AgNPs showed uniform spherical particles in the nano-range (about 20 nm). At the same time, TEM micrographs of the biosynthesized AgNPs revealed uniform nano particles and in the range of (10-25 nm). This result comes in agreement to some extent, with the finding of previous study conducted by El-Sherbiny et al., which showed that, the average particle diameters of myrrh AgNPs ranged from 40 to 100 nm. This particle size range was decreased to 10 to 50 nm by increasing the AgNO3 concentration to 6 mM followed by 60 min of UV irradiation (23).

In the current study, AgNPs showed antiparasitic activity whereas, ***T. spiralis*** larvae recorded high mortality rates after 2 days of exposure time at concentrations of 10, 15, 20 μg/ml. Compared with previous studies, the results agreed with Tomar and Preet, who revealed the lethal effect of AgNPs against ***Haemonchus contortus*** adult worm at different concentrations ranged from 1–25 μg/ml (33). Moreover, AgNPs significantly decrease oocyst viability of ***Cryptosporidium parvum*** in a dose-dependent manner, between concentrations of 0.005 and 500 μg/ml (34). Additionally, AgNPs at a concentration of 1 ppm for 30 min and 0.1 ppm for 1 h reduced the viability of ***Cryptosporidium*** oocysts by 97.2 and 94.4%, respectively (35).

Myrrh-biosynthesized AgNPs showed the highest lethal effect after 2 days of exposure at 5, 10, 15, 20 μg/ml. these results were in harmony with Barbosa et al., who showed that the green synthesis of AgNPs from ***nematophagous*** fungus exhibited a lethal effect on the infecting 3^rd^ stage larvae of ***Ancylostoma caninum*** (36). In the same way Rahimi et al., observed that, AgNPs derived from the aqueous aerial extract of ***Penicillium aculeatum*** had high scolicidal effects against protoscolices of hydatid cyst concentrations of 0.1 and 0.15 mg/mL after 120 min of exposure(37). The high mortality rate of larvae following AgNPs treatment could be attributed to the extremely small size of the nanoparticles, (10-25 nm), which probably permitted transcuticular absorption of AgNPs by the parasite causing the death of muscular larvae (36).

One of the hallmark effects of any anthelmintic agents was the destruction of the worm’s surface (38). In the current study, destruction of the cuticle of ***Trichinella*** muscular larvae by AgNPs and myrrh-biosynthesized AgNPs was obvious in the form of loss of striation, multiple vesicles, pores, sloughing of some areas and blebs on the cuticle. These findings were in agreement with the results of the previous study, which recorded the same forms of the cuticle destruction of ***T. spiralis*** muscular larvae when treated with AgNPs (39). Furthermore, Othman and Shoheib, revealed degenerative changes in the body wall of larvae and adults of ***T. spiralis*** under the influence of geldanamycin (40). Also, several studies were carried out to detect the effect of AgNPs on different parasites in vitro and reported that ***Schistosoma mansoni*** cercariae treated with AgNPs showed thinning of tegument with focal loss of spines and edematous swelling of the muscle layer (41). Gherbawy et al. observed perforations on the egg surface of ***Fasciola*** on *in vitro* exposure to AgNPs. The green synthesis of AgNPs from ***nematophagous*** fungus exhibited cuticle destruction on the third stage larvae of ***Ancylostoma caninum*** (36). The blebbing is an attempt by the parasite to replace damaged surface membrane in response to drug action (39). Silver nano-particles caused destruction of the cuticles through binding of the released positive silver ions to negatively charged cell membrane and interfere with membrane integrity (43). Silver ions may also adhere to the membrane wall, causing holes through which they also can penetrate inside the organism (44).

Chemically synthesized and Myrrh-biosynthesized AgNPs nearly exhibit the same influence on the ability of the treated ML to cause infection. They cause 100% reduction in adult and larval stages development in infected mice at sub-lethal dose. These results come in line with Bolás-Fernandez, who studied the ability of ***Trichinella spiralis*** muscular larvae to survive in different cell culture media and showed the dramatical reduction in larval infectivity when incubated under 5% CO2 and microaerobic conditions (27). Also, these findings agreed with the results of the study conducted by (45) who recorded complete block of cercarial infectivity on exposure of ***S. japonicum*** cercariae to AgNPs at conc. of 120µg/ml for 30 min. Liao et al., revealed that treatment of ***C. parvum*** oocysts with AgNPs prevent the infection outbreak (46). Su et al., used *in vitro* excystation assays to examine disinfection capabilities of AgNPs on ***C. parvum*** oocysts and validated their results by in vivo animal infectivity assays. A marked reduction in oocyst shedding among mice infected with AgNPs treated oocyst in comparison to controls (47).

In conclusion, this study revealed that AgNPs possess marked antiparasitic activity on ***T. spiralis*** larvae. Also, the current study verified that the Myrrh extract can be used as green reducing and capping agent for the biosynthesis of AgNPs with less hazardous effects on the environment. The developed AgNPs showed good stability with visible changes were observed even after 6 months. Moreover, the larvicidal effectiveness of chemically and green synthesized AgNPs against ***Trichinella*** larvae is almost the same with no significant difference, however, biosynthesis approach is more favorable because of its safety to humans and environment and easier method of preparation.

## Authors contribution

### Maintenance of *Trichinella spiralis* life cycle

Abeer El-sayed Mahmoud and Salwa Mahmoud Abd-ELrahman

### Methodology

Salwa Mahmoud Abd-ELrahman, Fahd M. Alsharif, Shaymaa M. Mohamed and Mosleh Mohammed Abomughaid

### Data analysis

Nahed Ahmed Elossily

### Supervision

Ahmed Kamal Dyab and Nahed Ahmed Elossily

### Writing – original draft

Abeer El-sayed Mahmoud, Nahed Ahmed Elossily and Salwa Mahmoud Abd-ELrahman

### Writing – review and editing

Ahmed Kamal Dyab Abeer El-sayed Mahmoud and Nahed Ahmed Elossily

## References

1. Yadav AK, Temjenmongla. Efficacy of Lasia spinosa leaf extract in treating mice infected with Trichinella spiralis. Parasitol Res. 2012;110(1):493–8.

2. Pozio E. World distribution of Trichinella spp. infections in animals and humans. Vet Parasitol. 2007;149(1–2):3–21.

3. Gottstein B, Pozio E, Nöckler K. Epidemiology, diagnosis, treatment, and control of trichinellosis. Clin Microbiol Rev. 2009;22(1):127–45.

4. Codina A V, García A, Leonardi D, Vasconi MD, Di Masso RJ, Lamas MC, et al. Efficacy of albendazole:β-cyclodextrin citrate in the parenteral stage of Trichinella spiralis infection. Int J Biol Macromol. 2015;77:203–6.

5. Shalaby MA, Moghazy FM. Effect of methanolic extract of Balanites aegyptiaca fruits on enteral and parenteral stages of Trichinella spiralis in rats Effect of methanolic extract of Balanites aegyptiaca fruits on enteral and parenteral stages of Trichinella spiralis in rats. 2014;(March 2010).

6. Basyoni MMA, El-Sabaa AAA. Therapeutic potential of myrrh and ivermectin against experimental Trichinella spiralis infection in mice. Korean J Parasitol. 2013;51(3):297–304.

7. Miśta D, Piekarska J, Houszka M, Zawadzki W, Gorczykowski M. The influence of orally administered short chain fatty acids on intestinal histopathological changes and intensity of Trichinella spiralis infection in mice. Vet Med (Praha). 2010 Jan 1;55(6):264–74.

8. Saxena A, Tripathi RM, Zafar F, Singh P. Green synthesis of silver nanoparticles using aqueous solution of Ficus benghalensis leaf extract and characterization of their antibacterial activity. Mater Lett [Internet]. 2012;67(1):91–4. Available from: http://dx.doi.org/10.1016/j.matlet.2011.09.038

9. Kaviya S, Santhanalakshmi J, Viswanathan B, Muthumary J, Srinivasan K. Biosynthesis of silver nanoparticles using citrus sinensis peel extract and its antibacterial activity. Spectrochim Acta - Part A Mol Biomol Spectrosc [Internet]. 2011;79(3):594–8. Available from: http://dx.doi.org/10.1016/j.saa.2011.03.040

10. Klaine SJ, Alvarez PJJ, Batley GE, Fernandes TF, Handy RD, Lyon DY, et al. Nanomaterials in the environment: Behavior, fate, bioavailability, and effects. Environ Toxicol Chem [Internet]. 2008 Sep 1;27(9):1825–51. Available from: https://doi.org/10.1897/08-090.1

11. Tsuji T, Kakita T, Tsuji M. Preparation of nano-size particles of silver with femtosecond laser ablation in water. Appl Surf Sci. 2003 Feb 1;206:314–20.

12. Lukman AI, Gong B, Marjo CE, Roessner U, Harris AT. Facile synthesis, stabilization, and anti-bacterial performance of discrete Ag nanoparticles using Medicago sativa seed exudates. J Colloid Interface Sci. 2011;353(2):433–44.

13. Durán N, Marcato PD, Alves OL, GIH De Souza, Esposito E. Mechanistic aspects of biosynthesis of silver nanoparticles by several Fusarium oxysporum strains. J Nanobiotechnology [Internet]. 2005;3(1):8. Available from: https://doi.org/10.1186/1477-3155-3-8

14. Kowshik M, Ashtaputre S, Kharrazi S, Vogel W, Urban J, Kulkarni S, et al. Extracellular synthesis of silver nanoparticles by a silver-tolerant yeast strain MKY3. Vol. 14, Nanotechnology. 2003. 95–100 p.

15. Justin Packia Jacob S, Finub JS, Narayanan A. Synthesis of silver nanoparticles using Piper longum leaf extracts and its cytotoxic activity against Hep-2 cell line. Colloids Surfaces B Biointerfaces. 2012;91:212–4.

16. Marimuthu S, Elango G, Kirthi AV, Jayaseelan C, Rajakumar G, Santhoshkumar T, et al. Evaluation of green synthesized silver nanoparticles against parasites. Parasitol Res. 2010;108(6):1541–9.

17. Said DE, ElSamad LM, Gohar YM. Validity of silver, chitosan, and curcumin nanoparticles as anti-Giardia agents. Parasitol Res. 2012;111(2):545–54.

18. Kar PK, Murmu S, Saha S, Tandon V, Acharya K. Anthelmintic efficacy of gold nanoparticles derived from a phytopathogenic fungus, Nigrospora oryzae. PLoS One. 2014;9(1):1–9.

19. Karamustafa SD, Mansour N, Ankli A, Baser KHC, Bickle Q, Tasdemir D. In vitro effect of Myrrh extracts on the viability of Schistosoma mansoni larvae. Planta Med. 2011 Aug 1;77:1317.

20. Gamble HR. Detection of Trichinellosis in Pigs by Artificial Digestion and Enzyme Immunoassay. J Food Prot. 2016;59(3):295–8.

21. Wassom D.L. DDA and DTA. T. spiralis infections of inbred mice: Immunological specific responses 13 induced by different Trichinella isolates. Parasitology. 1988;47(2):283–7.

22. Evans WC, Evans D, Trease GE. Trease and Evans’ pharmacognosy. Edinburgh; New York: WB Saunders; 2002.

23. El-Sherbiny IM, Salih E, Reicha FM. Green synthesis of densely dispersed and stable silver nanoparticles using myrrh extract and evaluation of their antibacterial activity. J Nanostructure Chem. 2013;3(1):8.

24. Mavani K, Shah M. Synthesis of Silver Nanoparticles by using Sodium Borohydride as a Reducing Agent. 2013.

25. Shoeb M, Singh BR, Khan JA, Khan W, Singh BN, Singh HB, et al. ROS-dependent anticandidal activity of zinc oxide nanoparticles synthesized by using egg albumen as a biotemplate. Adv Nat Sci Nanosci Nanotechnol [Internet]. 2013;4(3):35015. Available from: http://dx.doi.org/10.1088/2043-6262/4/3/035015

26. Bughdadi FA. Ultrastractural studies on the parasitic worm Trichinella Spiralis J Taibah Univ Sci [Internet]. 2013;3(1):33–8. Available from: http://dx.doi.org/10.1016/S1658-3655(12)60018-1

27. Bolás-Fernandez F. Total Anaerobiosis During In Vitro Culture in Conventional Cell Culture Media is Required for Retaining Infectivity of <span class=“genus-species”>Trichinella spiralis</span> L1 Larvae. J Parasitol [Internet]. 2002 Aug;88(4):794–6.Availablefrom: https://doi.org/10.1645/0022-3395(2002)088[0794:TADIVC]2.0.CO

28. Ahmad A, Wei Y, Syed F, Khan S, Khan GM, Tahir K, et al. Isatis tinctoria mediated synthesis of amphotericin B-bound silver nanoparticles with enhanced photoinduced antileishmanial activity: A novel green approach. JPhotochem Photobiol B Biol [Internet]. 2016;161:17–24. Available from: http://www.sciencedirect.com/science/article/pii/S101113441630210X

29. Mulvaney P. Surface Plasmon Spectroscopy of Nanosized Metal Particles. Langmuir [Internet]. 1996 Jan 1;12(3):788–800. Available from: https://doi.org/10.1021/la9502711

30. Moussa S, Mostafa A, Abdel-Raouf N, Ibraheem I. Biosynthesis of silver and silver chloride nanoparticles by Parachlorella kessleri SAG 211-11 and evaluation of its nematicidal potential against the root-knot nematode; Meloidogyne incognita. Aust J Basic Appl Sci. 2016 Dec 1;10:354–64.

31. Moussa S, Abdel-Alim M, Abdel-Raouf N, Ibraheem I. Biosynthesis of silver chloride nanoparticles using the cyanobacterium Anabaena variabilis. Life Sci J. 2017 May 1;14:25–30.

32. Sathishkumar M, Sneha K, Won SW, Cho C-W, Kim S, Yun Y-S. Cinnamon zeylanicum bark extract and powder mediated green synthesis of nano-crystalline silver particles and its bactericidal activity. Colloids Surfaces B Biointerfaces [Internet]. 2009;73(2):332–8. Available from: http://www.sciencedirect.com/science/article/pii/S0927776509002410

33. Tomar RS, Preet S. Evaluation of anthelmintic activity of biologically synthesized silver nanoparticles against the gastrointestinal nematode, Haemonchus contortus. J Helminthol. 2017;91(4):454–61.

34. Cameron P, Gaiser BK, Bhandari B, Bartley PM, Katzer F, Bridle H. Silver Nanoparticles Decrease the Viability of Cryptosporidium parvum Oocysts. Appl Environ Microbiol [Internet]. 2015 Oct 23;82(2):431–7. Available from: https://www.ncbi.nlm.nih.gov/pubmed/26497464

35. Hassan D, Farghali M, Eldeek H, Gaber M, Elossily N, Ismail T. Antiprotozoal activity of silver nanoparticles against Cryptosporidium parvum oocysts: New insights on their feasibility as a water disinfectant. J Microbiol Methods [Internet]. 2019;165:105698. Available from: http://www.sciencedirect.com/science/article/pii/S0167701219304002

36. Barbosa ACMS, Costa Silva LP, Ferraz CM, Tobias FL, de Araújo JV, Loureiro B, et al. Nematicidal activity of silver nanoparticles from the fungus <em>Duddingtonia flagrans</em>. Int J Nanomedicine [Internet]. 2019 Apr 2 [cited 2019 Nov 24];Volume 14:2341–8. Available from: https://www.dovepress.com/nematicidal-activity-of-silver-nanoparticles-from-the-fungus-duddingto-peer-reviewed-article-IJN

37. Rahimi MT, Ahmadpour E, Rahimi Esboei B, Spotin A, Kohansal Koshki MH, Alizadeh A, et al. Scolicidal activity of biosynthesized silver nanoparticles against Echinococcus granulosus protoscolices. Int J Surg [Internet]. 2015;19:128–33. Available from: http://dx.doi.org/10.1016/j.ijsu.2015.05.043

38. Shuhua X, Binggui S, Chollet J, Utzinger J, Tanner M. TEGUMENTAL CHANGES IN ADULT SCHISTOSOMA MANSONI HARBORED IN MICE TREATED WITH ARTEMETHER. J Parasitol [Internet]. 2000 Oct 1;86(5):1125–32. Available from: https://doi.org/10.1645/0022-3395(2000)086[1125:TCIASM]2.0.CO

39. Manal A, El-Melegy Ghoneim N, Mohamed N, El-Dien N, Rizk M. Silver Nano Particles Improve the Therapeutic Effect of Mebendazole Treatment during the Muscular Phase of Experimental Trichinellosis. 2019.

40. Othman AA, Shoheib ZS. Detrimental effects of geldanamycin on adults and larvae of Trichinella spiralis. Helminthol. 2016;53(2):126–32.

41. Moustafa MA, Mossalem HS, Sarhan RM, Abdel-Rahman AA, Hassan EM. The potential effects of silver and gold nanoparticles as molluscicides and cercaricides on Schistosoma mansoni. Parasitol Res. 2018;117(12):3867–80.

42. Gherbawy YA, Shalaby IM, El-Sadek MSA, Elhariry HM, Abdelilah BA. The anti-fasciolasis properties of silver nanoparticles produced by Trichoderma harzianum and their improvement of the anti-fasciolasis drug Triclabendazole. Int J Mol Sci. 2013;14(11):21887–98.

43. Marambio-Jones C, Hoek EM V. A review of the antibacterial effects of silver nanomaterials and potential implications for human health and the environment. J Nanoparticle Res. 2010;12(5):1531–51.

44. Jahangirian H, Lemraski EG, Webster TJ, Rafiee-Moghaddam R, Abdollahi Y. A review of drug delivery systems based on nanotechnology and green chemistry: green nanomedicine. Int J Nanomedicine. 2017 Apr;12:2957–78.

45. Cheng Y, Chen X, Song W, Kong Z, Li P, Liu Y. Contribution of silver ions to the inhibition of infectivity of Schistosoma japonicum cercariae caused by silver nanoparticles. Parasitology. 2013 Jan 24;140:1–9.

46. Liao KT, Varhue W, Guerrant RL, Smith JA, Swami NS. A simple and rapid method for infectious waterborne disease monitoring using disposable PDMS microfluidic chip by dielectrophoresis. Proc 16th Int Conf Miniaturized Syst Chem Life Sci MicroTAS 2012. 2012;1951–3.

47. Su Y-H, Tsegaye M, Varhue W, Liao K-T, Abebe L, Smith J, et al. Quantitative dielectrophoretic tracking for characterization and separation of persistent subpopulations of Cryptosporidium parvum. Analyst. 2014 Jan 7;139(1):66–73.

